# The Infralimbic, but not the Prelimbic Cortex is needed for a Complex Olfactory Memory Task

**DOI:** 10.1101/2024.10.15.618554

**Authors:** Dahae J. Jun, Rebecca Shannon, Katherine Tschida, David M. Smith

**Author notes:** Corresponding author: David M. Smith Department of Psychology, Cornell University.

## Abstract

The medial prefrontal cortex (mPFC) plays a key role in memory and behavioral flexibility, and a growing body of evidence suggests that the prelimbic (PL) and infralimbic (IL) subregions contribute differently to these processes. Studies of fear conditioning and goal-directed learning suggest that the PL promotes behavioral responses and memory retrieval, while the IL inhibits them. Other studies have shown that the mPFC is engaged under conditions of high interference. This raises the possibility that the PL and IL play differing roles in resolving interference. To examine this, we first used chemogenetics (DREADDs) to suppress mPFC neuronal activity and tested subjects on a conditional discrimination task known to be sensitive to muscimol inactivation. After confirming the effectiveness of the DREADD procedures, we conducted a second experiment to examine the PL and IL roles in a high interference memory task. We trained rats on two consecutive sets of conflicting odor discrimination problems, A and B, followed by test sessions involving a mid-session switch between the problem sets. Controls repeatedly performed worse on Set A, suggesting that learning Set B inhibited the rats’ ability to retrieve Set A memories (i.e. retroactive interference). PL inactivation rats performed similarly to controls. However, IL inactivation rats did not show this effect, suggesting that the IL plays a critical role in suppressing the retrieval of previously acquired memories that may interfere with retrieval of more recent memories. These results suggest that the IL plays a critical role in memory control processes needed for resolving interference.

## 1. Introduction

The medial prefrontal cortex (mPFC) is known to be involved in executive control (Euston et al., 2012; Funahashi, 2001; Rossi et al., 2009) and it plays a critical role in memory retrieval (Bontempi et al., 1999; Miller & Cohen, 2001; Tomita et al., 1999; Yadav et al., 2022). The PFC is critical for a variety of cognitive tasks that require subjects to resolve conflicting rules and responses (Miller, 2000), a function commonly referred to as behavioral flexibility (Ragozzino et al., 1999; Ragozzino, 2007). Consistent with this idea, PFC damage impairs strategy shifting tasks in humans and rodents (Birrell & Brown, 2000; Demakis, 2003; Milner, 1963; Ragozzino et al., 1999; Rich & Shapiro, 2007; Stuss et al., 2000). Our previous studies of olfactory memory in rodents have found that the mPFC is necessary when the memory demands of the task produce high levels of interference, but not when interference is minimal, suggesting that the presence of interference may be the critical factor that drives PFC engagement (Peters et al, 2013; Peters and Smith, 2020).

An extensive literature indicates that the rodent mPFC is not homogenous (Hoover & Vertes, 2007). Instead, the prelimbic (PL) and infralimbic (IL) cortices appear to play distinct roles in memory processes. Studies of fear conditioning (Milad & Quirk, 2002; Quirk et al., 2000) and action-outcome learning (Corbit & Balleine, 2003) have suggested that the PL and IL support opposing processes. Based on these and other similar findings, Gourley & Taylor (2016) proposed a “PL-go/IL-stop” model of the mPFC role in complex behaviors (but see Moorman & Aston Jones, 2015). Specifically, in studies of fear conditioning and extinction, the PL has been shown to promote the retrieval of a fear memory (Courtin et al., 2014; Do-Monte et al., 2015; Laurent & Westbrook, 2009; Sierra-Mercado et al., 2006,) while the IL inhibits retrieval (Laurent & Westbrook, 2009; Morgan et al., 1993; Quirk et al., 2000; Sierra-Mercado et al., 2011). The subregions also appear to differentially modulate stimulus and context associations (George et al., 2023). Similar to findings in fear studies, the PL has been observed to drive cocaine seeking behavior, while the IL suppresses the behavior after extinction (Mesa et al., 2022; Moorman et al., 2015). Furthermore, a study on avoidance and reward seeking demonstrated distinct roles of the subregions such that IL inactivation broadly impaired active and inhibitory avoidance while PL inactivation disrupted only active avoidance (Capuzzo & Floresco, 2020).

In the present study, we examined the role of the mPFC and its PL and IL subregions in two high interference olfactory memory tasks. In the first experiment, we used chemogenetics (DREADDs, Designer Receptors Exclusively Activated by a Designer Drug) to suppress neuronal activity in both the PL and IL, and we tested subjects on a conditional odor discrimination task that had previously been shown to be sensitive to muscimol inactivation of the mPFC (Devito et al, 2010). After replicating the previous study and confirming that our DREADDs procedure was effective, we conducted a second experiment aimed at determining whether the PL and IL play differing roles in high interference odor memory. For this experiment, we trained rats on two conflicting odor discrimination problem sets and used DREADDs inactivation to examine the role of each subregion during high interference test sessions involving a mid-session switch between the two problem sets.

## 2. Method

### Experiment 1: Conditional discrimination task

#### Subjects and Surgery

The subjects were 12 adult Long-Evans rats (6 females, 6 males, Charles River Laboratories, Wilmington, MA). One female was excluded from the analysis due to poor perfusion and tissue quality. The rats were housed singly and maintained on a 12-hr light-dark cycle, food restricted to 80-85% of their ad libitum weight and given free access to water. Prior to training, the rats were anesthetized with isoflurane, placed in a stereotaxic device, the skull was exposed, and craniotomies were drilled for DREADDs virus infusion. An adeno-associated virus, pAAV-hSyn-hM4D(Gi)-mCherry (AAV8), viral titer 2 x 10^13^ vg/mL, was injected bilaterally into the PL cortex (AP 2.9mm, ML ± 0.6, DV -4.2) and the IL cortex (AP 2.9, ML ±0.6, DV -5.4) using a Hamilton syringe and microinjection pump for a total volume of 250 nL per injection site. For the control group, pAAV-hSyn-mCherry (AAV8), viral titer 2.6 x 10^13^ vg/ml, was injected at the same coordinates. The rats were given an antibiotic (5 mg/kg Baytril) and an analgesic (5 mg/kg Ketoprofen) just prior to surgery. Rats were allowed to recover for 7-10 days before beginning behavioral training. Temporary inactivation of the medial prefrontal cortex was induced by i.p. injection of DREADDs agonist clozapine N-oxide (CNO, 5 mg/kg) twenty minutes prior to the relevant training sessions. All experiments were conducted in compliance with guidelines established by the Cornell University Institutional Animal Care and Use Committee.

#### Behavioral Training Procedures

Prior to training, rats were acclimated to the apparatus and then trained to dig in cups of corncob cage bedding material for buried rewards (45 mg purified formula precision pellets, Bioserve, Inc., Frenchtown, NJ). The apparatus was a wooden box (48 cm wide x 81 cm long x 51 cm deep) with three compartments, a black side, a white side and a neutral (tan woodgrain) compartment in the middle, which was equipped with dividers that could be removed to allow rats to access the black or white compartments. After acclimation and shaping, the rats were trained to perform a conditional discrimination task. Each trial began with the rat in the neutral center compartment, the divider was removed to allow the rat to access either the black or white compartment, and the rat was presented with two cups containing odorized bedding material (heptanol and ethyl valerate; pure odorants mixed into 10 mL of mineral oil to create a partial vapor pressure of 1 Pa and mixed into 2 liters of bedding material and stored in airtight containers; Cleland et al., 2002). The same two odors were presented in separate cups on every trial, with the conditional rule that one odor predicted a buried reward in the black compartment while the other odor contained the reward in the white compartment (i.e., black X+/Y- and white X-/Y+). The assignment of the odorant valence within each compartment was counterbalanced across rats, and the left and right position of the cups was randomized across trials. A digging response was recorded if the rat displaced any of the bedding, except incidental displacement (e.g., stepping into the cup while walking over it). The rat was allowed to dig until the reward was retrieved, then returned to the center compartment for an intertrial interval (ITI) of approximately 15 seconds while the experimenter prepared the cups for the next trial.

Rats were trained on the conditional discrimination rule using a sequence of training steps. First, the rats were given blocks of 10 trials in the black compartment with the relevant discrimination rule (e.g., heptanol is rewarded but ethyl valerate is not), followed by 10 trials in the white compartment with the reversed discrimination rule (e.g., ethyl valerate is rewarded but heptanol is not). When errors were made, corrections to dig in both cups were allowed for the first 3 sessions but were not allowed for subsequent sessions. The ten trial block sessions in each compartment continued until the rat achieved a behavioral criterion of 85% correct choices on two consecutive sessions. The rats were then trained on alternating blocks of 5 trials until they achieved 85% correct. Finally, the rats were given training sessions consisting of 64 trials with the black and white compartments presented in a random sequence. After rats achieved 85% correct on the final stage of training, they were given test sessions with CNO injections or saline control injections. The CNO test sessions took place at least 5 weeks after surgery to allow time for expression of the DREADDs receptors.

### Experiment 2: Odor set shifting task

#### Subjects and Surgery

The subjects were 30 adult Long-Evans rats (16 females, 14 males). Six rats were excluded from the analysis due to inaccurate placement or overexpression of DREADDs receptors outside the target subregions. Surgery took place prior to training and was similar to experiment 1, except that the virus was selectively injected bilaterally into either the PL cortex (AP 2.8mm, ML ± 0.6, DV 4.6) or the IL cortex (AP 2.8, ML ±3.3, DV 6.6) using a microinjector (Nanoject III, Drummond Scientific, Broomall, PA). Injections targeting the IL cortex were performed at a 15-degree angle from the midline in order to avoid inadvertent spread of the virus along the injector track into the overlying PL cortex. For the control group, pAAV-hSyn-mCherry (AAV8), viral titer 2.6x10^13^ vg/ml, was injected bilaterally into the PL or IL cortex. We injected 300 nL per site (20 nL pulse every 20 seconds with 10 seconds between pulses). CNO test sessions took place at least 5 weeks after surgery to allow time for expression of the DREADDs receptors.

#### Behavioral Training Procedures

We adapted procedures previously used in our laboratory to test olfactory memory under high interference conditions (Butterly et al., 2012; Peters & Smith, 2020). All training was done in a white Plexiglass chamber (45 cm wide x 60 cm long x 40 cm deep). Other materials as well as the procedures for acclimation and dig training were the same as experiment 1. After the rats learned to reliably retrieve the buried rewards, they began training on the first of two odor discrimination problem sets, each of which contained eight odor pairs (16 individual odors).

Twenty-four pure odorants served as cues, prepared as in experiment 1: Propyl butyrate, Ethyl acetate, Anisole, Ethyl isovalerate, Furfuryl propionate, n-Butyl glycidyl ether, 1-Butanol, n-Amyl acetate, Ethyl butyrate, Propionic acid, Benzaldehyde, 1-Octanol, Methyl 2-furoate, Butyl butyrate, Cis-3-Hexenyl acetate, Heptanol, Ethyl valerate, 5-Methylfurfural, D-Limonene, Methyl Butyrate, 2-Phenylethanol, 2-Furyl methyl ketone, 1-Nonanol, and Butyl Pentanoate.

For each trial, the rat was presented with the two odors comprising one of the eight discrimination problems, with one of the odors always rewarded and the other not rewarded. The predictive value of the odors (rewarded or non-rewarded) was counterbalanced across subjects and their locations (left or right side of the chamber) were randomized across trials. The daily training sessions consisted of 64 trials (eight trials with each odor pair, presented in an unpredictable sequence). After reaching a criterion of 90% correct choices on two consecutive sessions on the first problem set, the rats were trained on a second problem set, Set B. Each odor pair in Set B consisted of a novel odor and an odor which had previously been presented in Set A (Fig. 2). This ensured that the rats could not adopt a strategy of simply approaching the novel odor (or avoiding the familiar odor) for each new odor pair. The rats were given daily training sessions on Set B until they reached a behavioral criterion of 90% correct choices on two consecutive sessions. While learning Set B, all rats were given an i.p. injection of saline to acclimate them to the injection procedures to be used during the subsequent test sessions.

**Fig 1.**
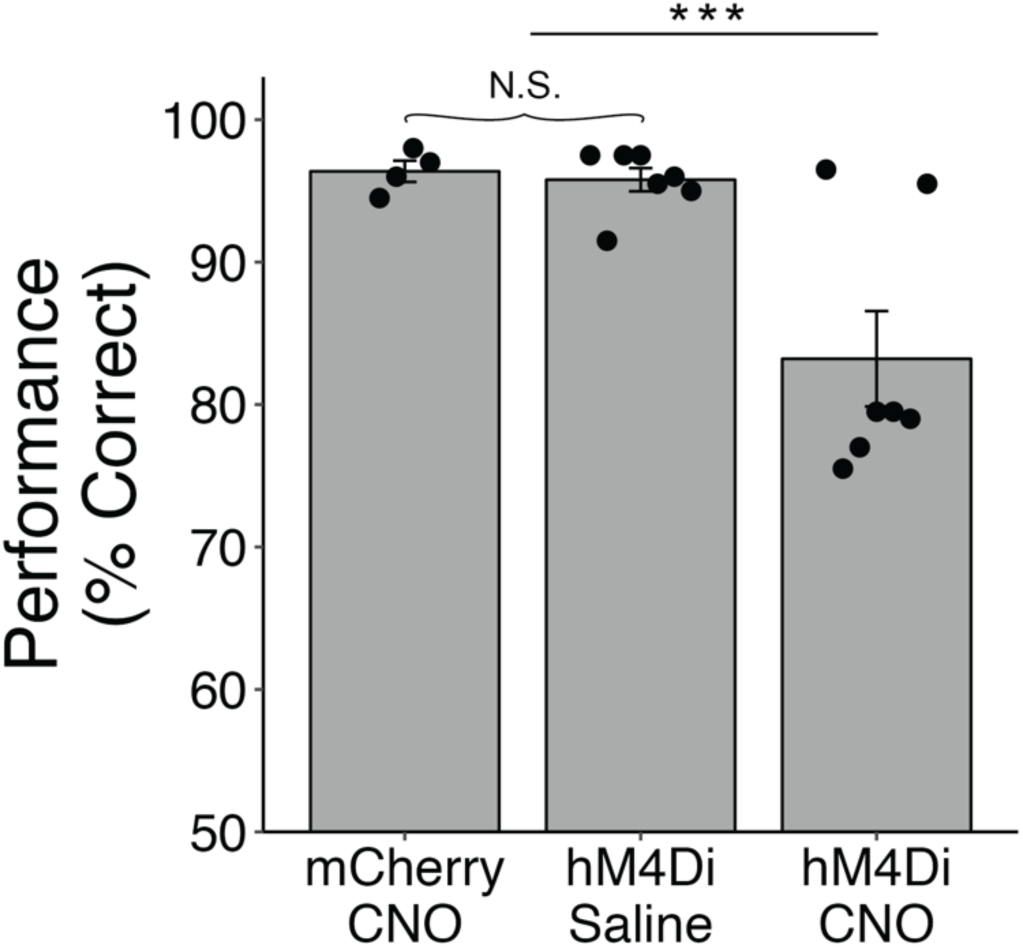
Performance of control subjects (mCherry CNO and hM4Di Saline) and DREADD inactivation subjects (hM4Di CNO) on the conditional discrimination task (*** indicates p < .001) with performance of individual subjects indicated by the dots.

**Fig 2.**
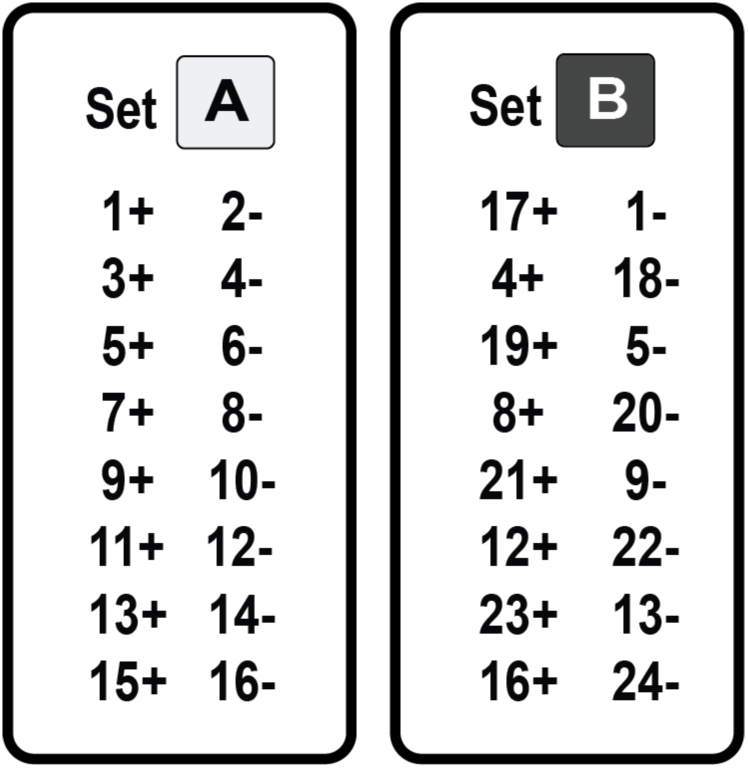
The odor discrimination problem sets presented in the odor set shifting task. Each number represents a distinct odor cue, and each problem set contains 16 odors (8 pairs). Set B consists of 8 novel odors and 8 familiar odors from Set A with their reward contingencies reversed.

After achieving the criterion for Set B, the rats were given three consecutive days of test sessions involving a mid-session switch between the two sets. The first half of each session (32 trials) was always the same as the problem set from the last half of the previous day’s session (Fig. 3A). For example, the first manipulation session consisted of 32 trials of Set B immediately followed by 32 trials of Set A. The mid-session switch from one problem set to the other was not cued. The second session started with 32 trials of Set A immediately followed by 32 trials of Set B, and so on for the last session.

**Fig 3.**
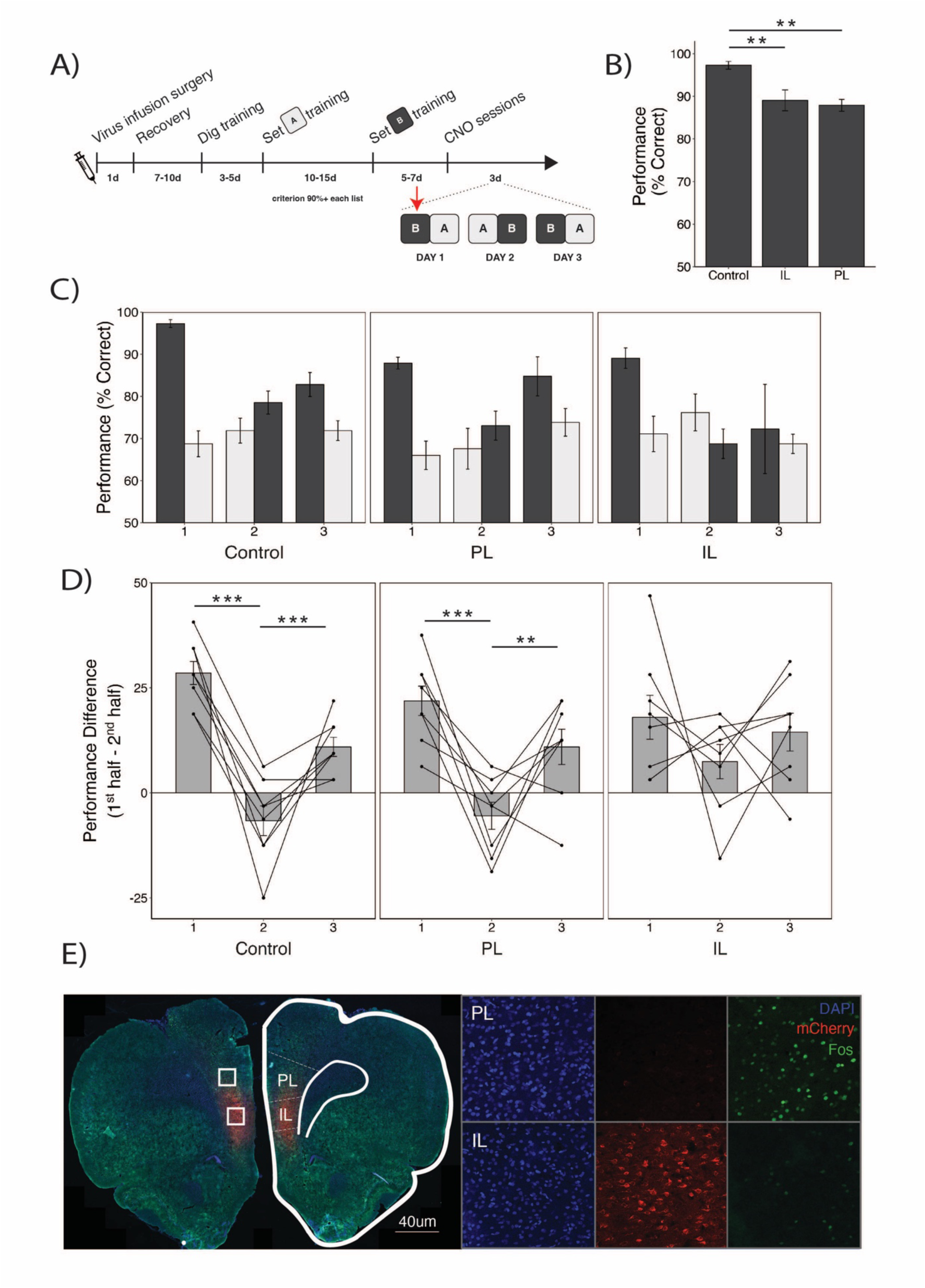
A) Experimental timeline for the odor set shifting task. B) Effects of CNO on ongoing performance on problem set B prior to any mid-session switch manipulations (red arrow). Inactivation of each subregion significantly impaired performance (*** indicates p < .001, ** p < 0.01). C) Performance of the three experimental groups during each of the three mid-session switch test sessions. Rats were given CNO injections prior to each of the test sessions. Performance is shown separately for the first and second halves of each test day, with problem set A shown in light grey and problem set B shown in black. Note that each test session began with the problem set from the previous day, as shown in A. D) Difference scores reflecting the change in performance (% correct) from the first half of each test session to the second half. The means (± SEM) are indicated by the bars, with the performance of individual subjects indicated by the dots and lines. E) Confocal image of a sagittal section from an example rat is shown, with hM4Di- mCherry expression targeted to the IL (red), DAPI (blue) and *c-Fos* (green). The rat was given CNO and 32 trials in the odor set shifting task. Note that *c-Fos* expression is apparent in the PL, where hM4Di was not expressed, but is largely absent in the IL. Scale bars = 50 um.

#### Perfusion and Tissue Processing

After the completion of the experiment, rats were deeply anesthetized with isoflurane and transcardially perfused with 0.1 M phosphate buffered saline (PBS) followed by 4% paraformaldehyde dissolved in 0.1 M PBS. Brains were extracted, post-fixed overnight in 4% paraformaldehyde dissolved in 0.1 M PBS before cryoprotection in 30% sucrose dissolved in PBS for 48 h before slicing. The brains were sectioned into 40-μm coronal slices, mounted on slides and stained with DAPI (ProLong Gold with DAPI). The brains of 2 control and 2 DREADDs rats were processed for cFos to visualize the effects of the inactivation procedures. Sections were washed 3 times in PBS for 5 minutes each, then incubated in 2% normal goat serum blocking solution with 0.1% Triton-X 100 for 1 hour at room temperature. The sections were then incubated in primary antibody (anti-c-Fos, #2250S, 1:2000, Cell Signaling, Danvers, USA) diluted in blocking solution overnight at room temperature. The following day, the sections were washed 3 times in PBS for 10 minutes each, then incubated in secondary antibody (Alexa Fluor 488 goat anti-Rabbit, 1:500, ThermoFisher, A-11008) at room temperature for 2 hours. After 3 more washes in PBS for 5 minutes each, the sections were mounted onto slides and cover-slipped with ProLong Gold Antifade Mountant with DNA Stains DAPI.

#### Data Analysis

Statistical analyses were performed using R. Two-way ANOVAs and Mixed-Factor ANOVA were computed as needed with factors of group, treatment, session, set, and set order. Tukey HSD *post hoc* tests were used to assess significance of differences between groups with alpha set to 0.05.

## 3. Results

### 3.1 Experiment 1: The mPFC Role in Conditional Discrimination

We assessed the effects of mPFC inactivation on the conditional odor discrimination task using a linear mixed effect model. Our control condition was comprised of hM4Di rats given saline injections (n = 7) as well as mCherry rats given CNO injections (n = 4). Therefore, we included in our model the fixed effects of group (hM4Di and mCherry) and treatment (saline and CNO) and random effect of rat ID. The two control conditions are shown separately in Figure 1. We found a significant interaction of the group and treatment conditions (F(1,9)=12.82, p = 0.006, Fig. 1). Post hoc (Tukey) comparisons confirmed that the two control groups were not different (p = 0.99), so we combined them into a single control group and found that the hM4Di CNO group performed significantly worse than controls (t(14.6) = 4.99, p = 0.0002, custom contrast used with Kenward-Roger approximation). Interestingly, two of the DREADDs subjects performed similarly to controls (Fig. 1). The reasons for this are unclear, as the hM4Di expression was not noticeably different for these subjects. Overall, the behavioral impairment is similar to a previous experiment using muscimol, suggesting that our chemogenetic approach was effective (also see Fig. 3E). We also examined differences in males and females and found no main effect of sex, and interactions of sex, group, and treatment were also not significant.

### 3.2 Experiment 2: The Role of PL and IL in a Proactive Interference Task

For this experiment, we trained rats on two conflicting odor discrimination problem sets (Fig. 2 and Method) and then gave them a series of three CNO test sessions involving a mid-session switch between the problem sets (Fig. 3A). Since the first CNO test session began with 32 trials of problem Set B (Fig. 3A, red arrow), the same problem set the rats had been performing for several days, our design offered the opportunity to determine whether PL or IL inactivation impaired ongoing performance on problem set B separately from the switch manipulation. We found that inactivation of either subregion significantly impaired performance (Fig. 3B). A two-way ANOVA on the data of problem set B on test day one with treatment group and sex as between-subject factors revealed a main effect of group (F(2,18) = 10.2, p = 0.001). Post hoc comparisons revealed that both PL (p = 0.002) and IL (p = 0.002) inactivated groups significantly differed from the control group but did not differ from each other (p = 0.99).

We then assessed the rats’ performance over the full three-day test sequence. Average performance on each test session and each half-session problem set is illustrated in figure 3C. In order to simplify the analysis and focus on the change in performance across the mid-session switch from one problem set to another, we computed a difference score between the first problem set of each day and the second (Fig. 3D), and we submitted these values to a to a mixed- factor ANOVA, with treatment group as a between-subjects factor (3 levels - Control, PL and IL) and session (3 levels - days 1-3) as a within-subjects factor. This analysis revealed a main effect of session (F(2,63) = 31.06; p < 0.0001) and a significant interaction of group and session variables (F(4,63) = 2.79; p = 0.03). We also examined differences in males and females and found no main effect of sex nor significant interactions of sex, group, and session; thus, we do not further discuss effect of sex. Post hoc comparison revealed a difference in the pattern of performance across sessions for the three groups. For both control subjects and subjects with PL inactivation, we observed a pattern in which the rats tended to perform better on set B than set A. Because the order of presentation of the odor sets varied across sessions, B-A, followed by A-B and then B-A (see Fig. 3A), this resulted in difference scores that were high for the first session, low for the second session and high again on the third session. In contrast, subjects with IL inactivation showed no such pattern. Instead, the rats tended to perform better on the first problem set of the day, regardless of whether that was Set A or Set B, resulting in positive difference scores which did not change significantly across the three sessions.

## 4. Discussion

In two experiments, we found that the mPFC plays an important role in olfactory memory processes. In our first experiment, we found that combined inactivation of the PL and IL impaired performance on a contextually-cued conditional discrimination task, consistent with previous studies (DeVito et al, 2010). Because the predictive value of the odor cues is reversed in the black and white contexts, subjects had to learn a complex conditional rule, manage conflicting response tendencies, and generally exhibit the kind of behavioral flexibility that is a hallmark of PFC functions (Euston et al., 2012; Navawongse & Eichenbaum, 2013; Ragozzino et al., 2003). However, there are also conflicting memory demands since subjects must remember which odor is associated with reward in the two contexts, so our result is also consistent with theoretical accounts suggesting that the PFC mediates cognitive control over memory retrieval processes (Bontempi et al., 1999; Corcoran & Quirk, 2007; Frankland et al., 2004; Takashima et al., 2006). One observation from our second experiment particularly supports this idea.

Previously, we found that mPFC inactivation impaired performance on a single odor discrimination problem set like those employed here (Peters et al., 2013). This impairment was likely attributed to the requirement that subjects simultaneously manage many odor memories, since there was no impairment when they were allowed to learn one discrimination problem at a time. In the present study, we found that inactivation of each individual subregion produced a modest, but statistically reliable impairment in ongoing performance on problem set B (Fig. 3B). This occurred before subjects were exposed to the mid-session switch manipulation, so the impairment could not be due to changing rules or response requirements (also see Peters and Smith, 2020).

In our second experiment, we tested subjects’ ability to perform a mid-session switch between two conflicting odor discrimination problem sets (Fig. 2). Unlike the conditional discrimination task, where rats could use the background context to determine which odor was rewarded on any given trial, there was no explicit cue to inform the rats about the current problem set. Instead, the rats had to deduce which set of rules was in effect and respond accordingly. Control subjects were readily able to do this, performing well above chance levels throughout the test sessions. This is consistent with numerous studies showing that intact rats are capable of set-shifting and rule-switch tasks (Birrell & Brown, 2000; Dias & Aggleton, 2000; Ragozzino et al., 1999, 2003, 2007; Rich & Shapiro, 2007). However, their performance was notably better for problem set B, the most recently learned of the two problem sets. This finding suggests that learning problem set B impaired memory for the previously learned problem set A (i.e. retroactive interference, Underwood, 1957). This effect was quite striking. Of the 21 test sessions conducted in seven rats over three days, performance was better on set B more than 90% of the time. This occurred despite strong initial learning of problem set A, with all the rats achieving greater than 90% correct, and this effect persisted throughout the three testing sessions even though the rats received 36 trials with problem set A each day. Rodents typically exhibit very strong and persistent odor memory (Tong et al., 2014; Wang et al., 2020), suggesting that the present results are not likely due to passive forgetting of problem set A. Instead, we suggest that this effect is the result of active suppression of problem set A memories that is caused by learning the conflicting memories of problem set B. An extensive body of work has shown that memory control processes mediated by the PFC can involve suppression of conflicting memories (Anderson et al., 1994; Anderson & Neely, 1996; Bekinschtein et al., 2018; Wimber et al., 2015; Wu et al., 2014;). In the case of our control subjects, the poorer performance on problem set A may have been mediated by the functioning of the intact PFC, particularly the IL cortex as we discuss below.

In contrast to control subjects, rats with IL inactivation did not show reliably better memory for problem set B. Instead, they tended to perform better on whichever problem set was presented first on each of the test days, regardless of whether it was problem set A or set B. This became apparent during test session two when, unlike controls, IL-inactivation rats performed better on problem set A (see light grey bar for Day 2 in Fig. 3C and positive values in Fig. 3D for IL rats, compared to Control and PL rats). In our experimental design, the first problem set for each test day was the same as the end of the previous day (Fig. 3A). Thus, IL inactivation resulted in better memory for the most recently experienced problem set on test day 2, without the apparent retrieval advantage of set B memories seen in controls. To the extent that intact controls experienced suppression of problem set A memories, as described above, IL inactivation appears to have blocked this memory suppression effect, suggesting that the IL plays an important role in suppressing conflicting memories.

Some models of PFC function suggest that the PL and IL play opposing roles in modulating memory retrieval processes. As discussed above, studies of fear conditioning and extinction demonstrate the differential roles of the PL and IL in promoting and inhibiting memory retrieval (Do-Monte et al., 2015; Otis et al., 2017; Quirk et al., 2000). Specifically, stimulation of PL neuronal activity increases fear retrieval (Vidal-Gonzalez et al., 2006) and PL inactivation reduces fear retrieval (Corcoran & Quirk, 2007; Laurent & Westbrook, 2009; Sierra-Mercado et al., 2011). Manipulation of IL neuronal activity has the opposite effect: inactivation increases retrieval (Morgan et al., 1993; Sierra-Mercado et al., 2006, 2011) while stimulation reduced retrieval (Burgos-Robles et al., 2007; Do-Monte et al., 2015; Vidal-Gonzalez et al., 2006).

Although this PL-go/IL-stop dichotomy is commonly cited in fear conditioning studies (see Gourley & Taylor, 2016), a growing literature in goal directed learning also supports this idea (Bari et al., 2011; Cholvin et al., 2016; Gutman et al., 2017; Ostlund & Balleine, 2005; Pfarr et al., 2015; Tran-Tu-Yen et al., 2009; Van Holstein & Floresco, 2020). In our study, the results of IL inactivation were consistent with this idea insofar as the loss of IL activity resulted in reduced inhibition of the set A memories.

According to this theoretical framework, PL inactivation might have been expected to reduce retrieval of either set A or set B memories, but we found no changes in performance during the mid-session switch sessions. The reasons for this are not clear. However, it is possible that our task prioritizes cognitive control processes that inhibit retrieval over those that promote retrieval. Because the rats received extensive training prior to the test sessions, the odor-reward associations of both problem sets were presumably very strong, and there may have been little need to promote the retrieval of these already-strong memories. Instead, the requirement for subjects to rapidly switch between the two strong, but conflicting sets of memories may have preferentially engaged retrieval inhibition processes, rendering PL neuronal activity irrelevant to performance.

This account positions the mPFC as a modulator of memory retrieval rather than a storage site for the odor memories themselves, and previous studies have shown that the mPFC is not needed for basic memory tasks that do not involve interference or complex rule switching (Birrell & Brown, 2000; Peters et al., 2013; Seamans et al., 1995). However, when interference does present a problem for subjects, the PL and IL are well-positioned to influence the olfactory regions of the brain where odor memories may be stored. In particular, these mPFC subregions have extensive anatomical projections to the anterior olfactory nucleus (Vertes, 2004).

Consistent with this idea, inactivation of the anterior olfactory nucleus impairs performance on the conditional discrimination task used in our first experiment (Levinson et al, 2020) and preliminary data from our laboratory show that neurons in this region respond to the odor cues and their valence in the odor set shifting task used for experiment two (Wu et al, 2023). Thus, complex olfactory memory tasks, such as those employed here, may be a particularly useful approach for examining the memory functions of the mPFC and its subregions.

## Acknowledgments

The authors would like to thank Sabrina Giaimo, Lizbeth Genao, and Aerin Mok for assistance with data collection. We would also like to thank Matt Thomas, Ph.D. at the Cornell Statistical Consulting Unit for assistance with data analysis.

## Funding

This work was supported by NIMH grant MH083809 and the College of Arts & Sciences New Frontier Grants to D. Smith.

## Declaration of interest

none

## Authors’ contribution

*D.J. Jun*: conceptualization, methodology, investigation, validation, formal analysis, writing – original draft, writing – review & editing, visualization

*R. Shannon*: investigation

*K. Tschida*: supervision

*D.M. Smith*: conceptualization, methodology, writing – review & editing, resources, supervision, funding acquisition

## Notes

### Competing Interest Statement

The authors have declared no competing interest.

